# EnzymeCAGE: A Geometric Foundation Model for Enzyme Retrieval with Evolutionary Insights

**DOI:** 10.1101/2024.12.15.628585

**Authors:** Yong Liu, Chenqing Hua, Tao Zeng, Jiahua Rao, Zhongyue Zhang, Ruibo Wu, Connor W Coley, Shuangjia Zheng

**Affiliations:** Global Institute of Future Technology, Shanghai Jiaotong University.; Department of Chemical and Biological Engineering, Hong Kong University of Science and Technology.; School of Pharmaceutical Sciences, Hainan University.; School of Computer Science and Engineering, Sun Yat-sen University.; School of Computer Science, McGill University.; Mila-Quebec AI Institute; School of Pharmaceutical Sciences, Sun Yat-sen University.; Department of Chemical Engineering, MIT.; Department of Electrical Engineering and Computer Science, MIT.

## Abstract

Enzyme catalysis is fundamental to life, driving the chemical transformations that sustain biological processes and support industrial applications. However, unraveling the intertwined relationships between enzymes and their catalytic reactions remains a significant challenge. Here, we present EnzymeCAGE, a catalytic-specific geometric foundation model trained on approximately 1 million structure-informed enzyme-reaction pairs, spanning over 2,000 species and encompassing an extensive diversity of genomic and metabolic information. EnzymeCAGE features a geometry-aware multi-modal architecture coupled with an evolutionary information integration module, enabling it to effectively model the nuanced relationships between enzyme structure, catalytic function, and reaction specificity. EnzymeCAGE supports both experimental and predicted enzyme structures and is applicable across diverse enzyme families, accommodating a broad range of metabolites and reaction types. Extensive evaluations demonstrate EnzymeCAGE’s state-of-the-art performance in enzyme function prediction, reaction de-orphaning, catalytic site identification, and biosynthetic pathway reconstruction. These results highlight its potential as a transformative foundation model for understanding enzyme catalysis and accelerating the discovery of novel biocatalysts.

## 1 Main

Enzymatic catalysis is fundamental to the complex biochemical networks essential for life, driving diverse metabolic processes with remarkable specificity and efficiency [1–4], and plays an essential role in a wide range of industries, from pharmaceuticals to biofuels [5–8]. Understanding and predicting enzyme functions is important to advancing both fundamental biology and practical applications. In this context, enzyme function prediction refers to two key capabilities: (1) predicting the functions of uncharacterized enzymes within a list of known enzymatically-catalyzed reactions, aiming to identify new or more efficient biocatalysts; (2) specifying a desired reaction and then virtually retrieving enzyme sequences/structures for compatibility. These two capabilities are central to understanding enzymatic roles, driving the discovery of novel biocatalysts and fostering innovations in synthetic biology.

However, accurately and systematically predicting enzyme functions remains a major challenge in biotechnology [9–12]. This challenge is underscored by significant annotation gaps in current databases: among the roughly 190 million protein sequences cataloged in UniProt [13], fewer than 0.3% are curated by experts, with only 19.4% of these supported by experimental evidence. Meanwhile, metabolic databases such as KEGG and MetaCyc [14, 15] report that 40-50% of known enzymatic reactions lack associated enzyme sequences, termed “orphan” reactions. These gaps in knowledge constrain our understanding of metabolic networks and limit the development of synthetic biology and metabolic engineering applications [16–18].

To bridge these gaps, numerous computational methods have been developed to link enzymes with the reactions they catalyze [19–26]. One approach to partially address this task involves mapping enzymes to Enzyme Commission (EC) classes—a hierarchical system that categorizes enzyme functions by four-digit codes [27–32]. However, a single EC class can represent multiple distinct reactions and novel reactions may not fit existing classifications [33–35]. Other approaches focus on directly predicting enzyme-substrate interactions [23, 24, 36–39], often employing similarity-based methods like Selenzyme [40] and BLASTp [41]. Their dependence on sequence similarity can lead to unreliable predictions, especially for enzymes with low sequence homology [42, 43].

As a result, there remains an urgent need for more advanced computational tools that can provide accurate and comprehensive functional predictions, ultimately enabling the discovery of novel enzyme functions and a deeper understanding of orphan reactions.

To address these needs, we introduce **EnzymeCAGE**, a **CA**talyticspecified **G**eometric-**E**nhanced **Enzyme**-reaction foundation framework that connects enzymes and reactions by integrating enzyme structures, evolutionary insights, and reaction center transformations. Unlike traditional EC-based methods, EnzymeCAGE uses a geometry-aware, reaction-guided interaction module to model the viability of an enzyme to catalyze a given reaction directly. This module identifies catalytic regions within an enzyme using a pocket-based attention mechanism, enhancing the interpretability and robustness of the model’s predictions. By pinpointing active regions, the model offers more interpretable and reliable predictions than traditional alignment-based or machine learning-based approaches. EnzymeCAGE is trained and evaluated using a genome-scale dataset comprising 2962 species to establish a robust foundation for understanding catalytic systems across various taxa. Our empirical results demonstrate that EnzymeCAGE excels in anticipating the function of unseen enzymes and annotating orphan reactions with the enzymes that catalyze them, demonstrating better retrieval results through a sequence similarity-free retrieval approach that is orthogonal to conventional methods. EnzymeCAGE quantitatively demonstrates robust extrapolation on unseen enzymes and orphan reactions, achieving a 44% improvement in enzyme function prediction and a 73% increase in retrieving enzymes for orphan reactions. Through simple fine-tuning, EnzymeCAGE can quickly adapt to family-specific tasks, improving its predictive accuracy within specific enzyme families. To illustrate its practical utility, we present a case study involving glutarate synthesis, where EnzymeCAGE accurately reconstructs a natural product pathway and significantly outperforms current state-of-the-arts methods. These findings highlight EnzymeCAGE’s potential as a powerful tool for enzyme retrieval, function prediction, and pathway engineering.

## 2 Results

### 2.1 EnzymeCAGE Approach

We illustrate the EnzymeCAGE approach in Fig. 1, where it takes an enzyme structure and a reaction as input and returns an overall catalytic score for their compatibility. EnzymeCAGE begins by extracting catalytic pockets using AlphaFill [44], then encodes geometric features of the pocket, including backbone atomic coordinates, dihedral angles, and side-chain torsions. This pocket region is represented as a graph: nodes correspond to residues with features such as residue type and geometric encodings, while edges represent bonds and residue connections, with features including atomic distances and orientations. We employ a Graph Neural Network (GNN) to encode the local catalytic pocket structures. EnzymeCAGE also incorporates embeddings from a protein language model ESM2 [45] to provide a more comprehensive representation that combines both local pocket-level and global enzyme-level features to describe the enzyme’s catalytic potential. To encode the reaction, EnzymeCAGE first computes substrate and product conformations, then identifies the reaction center that represents the subset of atoms undergoing changes in covalent bonding. Atoms within the reaction center are assigned higher weights to emphasize their relevance. We use a molecular GNN to encode the conformations of both substrate and product. Further methodological details are discussed in the Methods section.

**Fig. 1.**
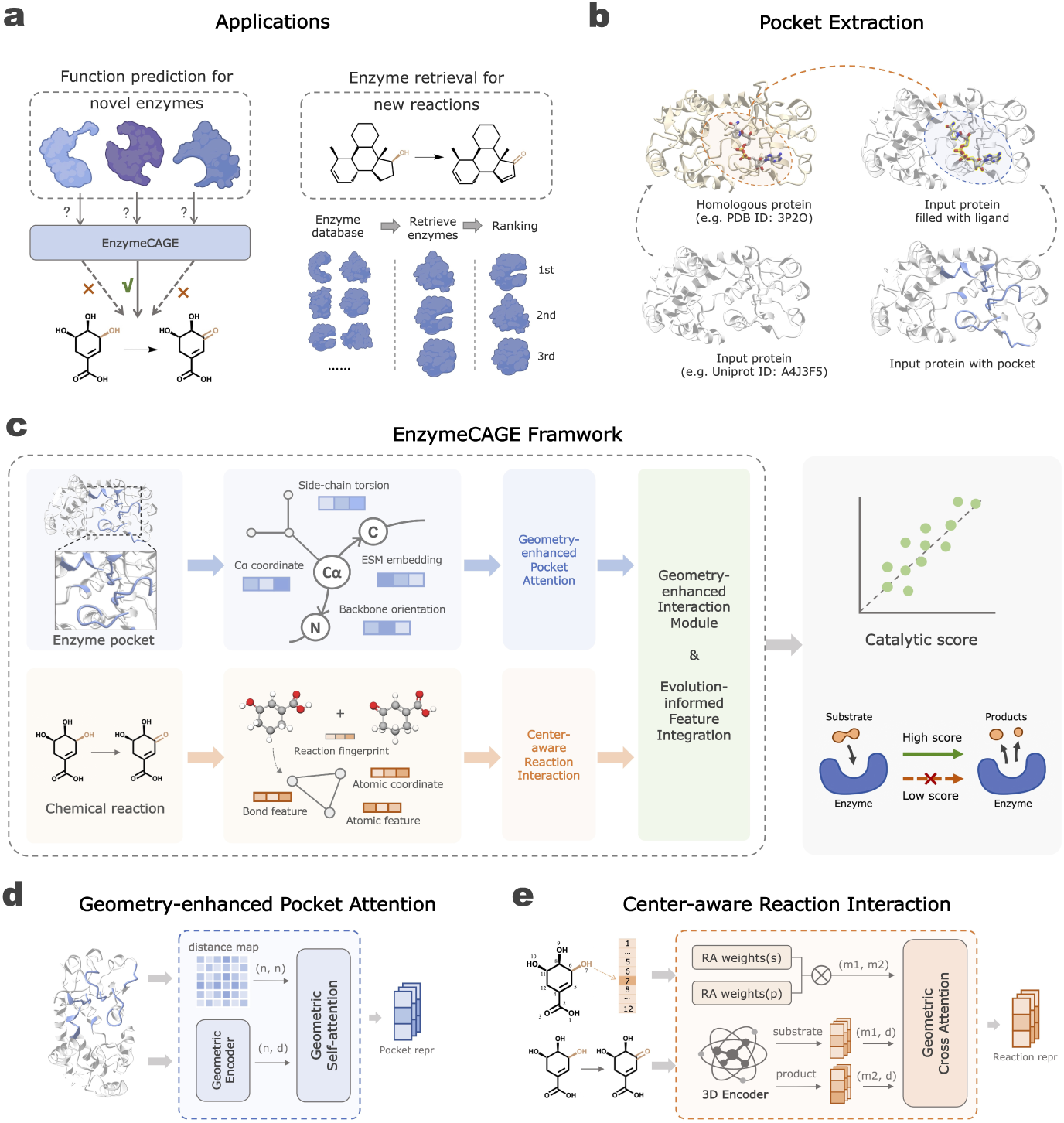
**Overview of EnzymeCAGE**. **(a)** Applications of EnzymeCAGE: function prediction for unseen enzymes and enzyme retrieval for new reactions. **(b)**Pocket extraction by AlphaFill. **(c)** EnzymeCAGE is a deep learning framework designed to predict enzyme-reaction catalytic specificity by encoding both pocket-specific enzyme structures and chemical reactions. **(d)** The Geometry-enhanced Pocket Attention Module uses a distance map between residues within the catalytic pocket as geometric guidance, which is then fed into a self-attention model to encode relative positional information among residues. **(e)** The Center-aware Reaction Interaction Module identifies reaction center, creating reacting area weights for both substrate and product atoms. The outer product of these weights generates a reacting area weight matrix, which is input as geometric guidance into a cross-attention model.

EnzymeCAGE further processes these representations of the enzyme pocket and chemical reaction in two separate network trunks before modeling their interactions. First, it uses a geometry-enhanced pocket attention module, constructing a distance map between pocket residues as an attention bias in a self-attention block to incorporate the pocket’s spatial structure. Concurrently, a center-aware reaction interaction module captures dynamic substrate-product changes, using cross-attention guided by reaction-center weights. The resulting pocket and reaction embeddings are combined using a geometry-enhanced interaction module that computes 3D interaction features between pocket residues and reaction molecules. These interaction features are integrated with ESM-derived enzyme features and DRFP [46] fingerprints of the reaction and fed into a final output network to predict catalytic activity— a numerical score normalized between 0 and 1 that describes the enzyme’s capability to catalyze the given reaction.

#### Dataset and Model Training

Foundation models like EnzymeCAGE differ from previous locally trained models. Instead of developing separate models for each enzyme family, EnzymeCAGE is a unified pretrained model that can adapt to various enzyme families and reactions. Here, we compiled a large-scale dataset of curated enzyme-reaction pairs from RHEA [14], MetaCyc [15], and BRENDA [47]. To optimize the model training process, we preprocessed the dataset by refining the protein and reaction data, removing invalid entries, and standardizing the data format. Additionally, we employed two novel strategies to generate diverse and high-quality negative samples based on enzyme functional similarity and reaction template perspectives. To conclude, we constructed a dataset comprising over 914,077 enzyme-reaction pairs (more details discussed in Methods). To evaluate model performance on unseen enzymes and novel reactions, we constructed two test sets, ”Loyal1968” and ”Orphan-194”, and created corresponding training sets to ensure proper train/test separation, as detailed in the Methods.

To train EnzymeCAGE, we constructed specific training sets for each evaluation scenario: (a) For the Loyal-1968 evaluation set, we excluded enzyme-reaction pairs present in Loyal-1968 from the training set. (b) For the Orphan-194 evaluation set, we included only reactions annotated with EC numbers and their associated data in the training set. EnzymeCAGE was trained end-to-end using binary cross-entropy loss as the loss function and Adam as the optimizer [48]. Overall, EnzymeCAGE showed better performance in enzyme retrieval and function prediction, surpassing state-of-the-art methods in accuracy, interpretability, and generalizability, even when using full enzyme structures. In addition, we conducted ablation experiments to verify the contribution of each module in EnzymeCAGE(Fig. 2g).

**Fig. 2.**
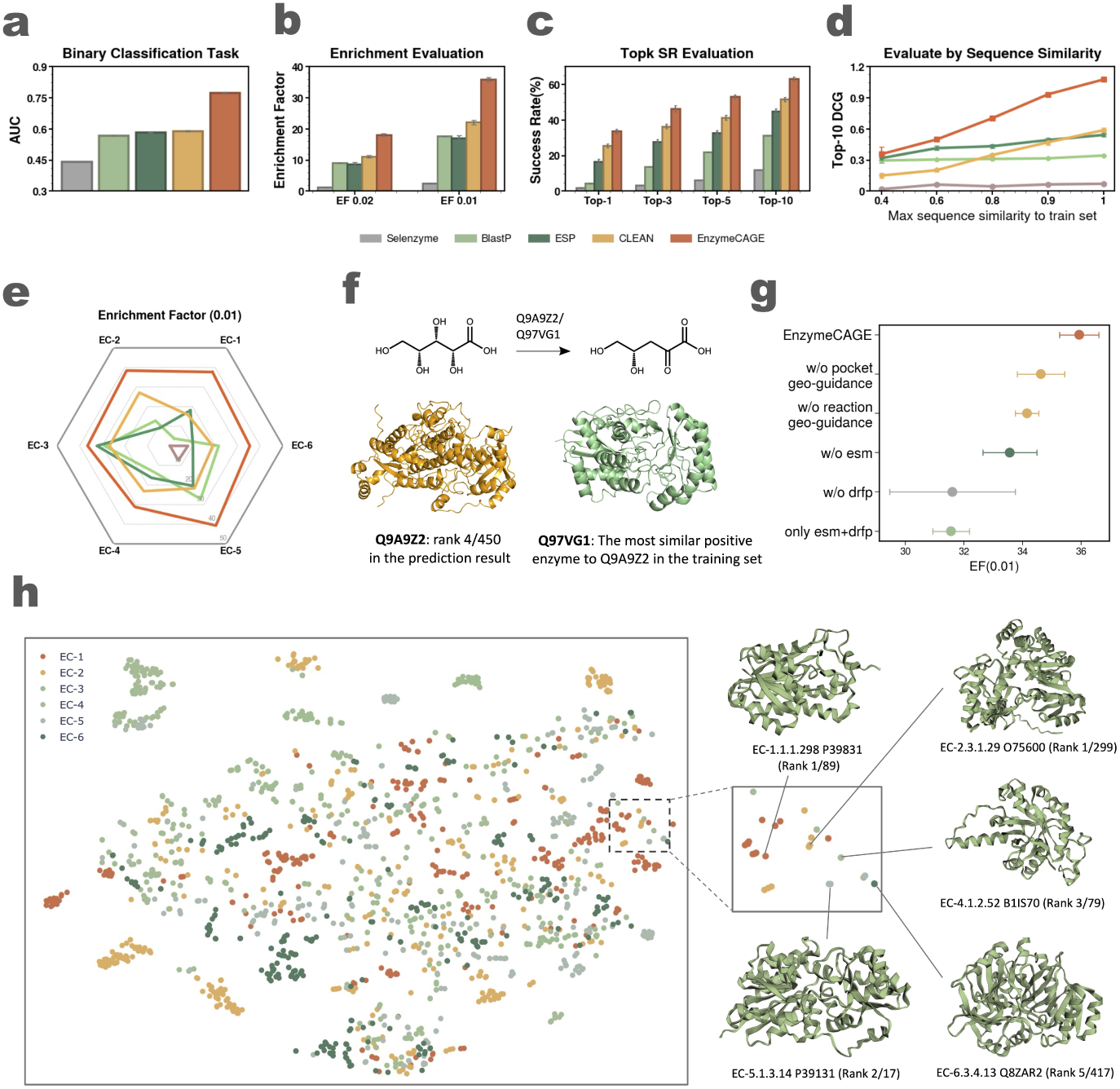
**EnzymeCAGE’s performance on test set Loyal-1968**. **(a)** Prediction performance comparison between EnzymeCAGE and baseline methods on the binary classification task. **(b)** Retrieval results evaluated using enrichment evaluation at two ratios of 0.01 and 0.02. **(c)** Top-k Success Rate (SR) evaluation results. **(d)** Top-10 Discounted Cumulative Gain (DCG) evaluation across varying thresholds of maximum sequence similarity to the training set. **(e)** Results after dividing reactions based on different EC classifications. **(f)** Example demonstrating EnzymeCAGE’s ability to accurately predict functionally similar enzymes with substantially different structures. Enzyme Q97VG1 (green) and Q9A9Z2 (yellow) catalyze the same reaction. Q9A9Z2, included in the test set, has Q97VG1 as its most structurally similar enzyme in the training set, with an alignment RMSD of 18.45. EnzymeCAGE successfully ranks Q9A9Z2 as a positive enzyme, placing it 4th out of 450 candidates. **(g)** Ablation results using Enrichment Factor (EF) as the evaluation metric. **(h)** t-SNE visualization of the positive sample enzyme distribution based on ESM features. The right panel provides an example showcasing EnzymeCAGE’s capacity to distinguish between enzymes with similar sequences but distinct functions. In this selected region, five enzymes in the test set Loyal-1968 were chosen based on sequence similarity but catalyze different reactions. EnzymeCAGE correctly identifies the positive enzyme for each reaction, yielding top-ranking results for each.

### 2.2 EnzymeCAGE Peformance on Enzyme Function Prediction

We first assessed EnzymeCAGE’s ability to predict enzyme functions by evaluating the model on unseen enzymes. We constructed a diverse test set, named ”Loyal-1968”, consisting of seen reactions and 1968 non-promiscuous(unseen) enzymes. In Loyal-1968, positive samples include non-promiscuous enzymes with a broad functional range across 77 unique top-three-level EC numbers, spanning 522 species, while negative samples consist of homologous enzymes with different functions. We provide more details on the construction of Loyal1968 in the Methods section. Loyal-1968 represents a real-world application scenario for enzyme function prediction, where an enzyme with unknown function may exhibit high sequence similarity to annotated enzymes but have different functional capabilities.

To evaluate EnzymeCAGE’s performance in enzyme function prediction within the Loyal-1968 test set, we examined its ability to rank positive enzymes prominently in a retrieval list for each reaction. EnzymeCAGE was benchmarked against state-of-the-art enzyme function prediction methods, including two similarity-based approaches, Selenzyme [40] and BLASTp [41], as well as two deep learning-based methods: ESP [24], CLEAN [32]. Four evaluation metrics were used to assess retrieval performance: Area under the curve (AUC) [49], Top-10 Discounted Cumulative Gain (DCG) [50], Top-k Success Rate (SR) [26], and Enrichment Factor (EF) [51].

For Top-k SR, EnzymeCAGE achieved a 33.7% Top-1 accuracy and a 63.2% Top-10 accuracy for function prediction (Fig. 2c), outperforming baselines by significant margins. These results indicate EnzymeCAGE’s better accuracy and success rate in function prediction. Additionally, EnzymeCAGE outperformed all baselines on binary classification AUC (Fig. 2a), EF (Fig. 2b), Top-k SR (Fig. 2c), and Top-10 DCG (Fig. 2d) evaluations, highlighting not only improved function prediction results but also a greater enrichment of true enzymes among high-scoring predictions. When evaluated by EC class, EnzymeCAGE demonstrated better performance across four out of six EC classes (EC-2, EC-3, EC-5, EC-6) with higher EF scores (Fig. 2e).

EnzymeCAGE also excelled in specific case studies of enzyme function prediction. In the first case, EnzymeCAGE successfully identified an unseen enzyme-reaction pair (Fig. 2f), ranking the correct enzyme (UniProt: Q9A9Z2, in yellow) in the 4th position among 450 candidates, despite substantial structural divergence from the most similar enzyme (UniProt: Q97VG1, in green) in the training set. This highlights EnzymeCAGE’s robustness in recognizing functional relevance in structurally diverse enzymes. In the second case, EnzymeCAGE distinguished between enzymes with high sequence similarity but distinct functions (Fig. 2h). For 1,968 positive enzymes in the test set, ESM2 embeddings and tSNE [52] were used to visualize the embeddings, selecting a cluster of five enzymes with different EC classes, indicating high sequence similarity yet functional diversity. EnzymeCAGE successfully retrieved all five enzymes in top positions within their candidate lists, emphasizing its capacity to discern functionally distinct enzymes despite sequence similarity.

Beyond function prediction for unseen enzyme-reaction pairs, EnzymeCAGE can pinpoint catalytic regions for unseen enzymes (Supplementary Fig. S4)—a capability that is neither a direct training objective nor derived from annotated training data. To validate this, we first gathered enzymes from the Catalytic Site Atlas [53], a database containing experimentally validated annotations of catalytic pockets. During predictions for these enzymes, we extracted attention weights from the enzyme-reaction interaction module. Each entry in the attention matrix reflects residue-atom interactions between enzyme residues and substrate atoms. We then focused on columns related to the substrate’s reaction center, computed per-residue mean values, and normalized the resulting weight vector. This allowed us to assess if these residues were part of the true catalytic pockets. EnzymeCAGE successfully highlighted catalytic regions across various enzymes, as validated against known annotations.

### 2.3 EnzymeCAGE Performance on Reaction De-orphaning

We further assessed EnzymeCAGE’s ability to capture catalytic specificity by evaluating its enzyme retrieval performance on orphan reactions. Here, EnzymeCAGE aims to identify and retrieve suitable candidate enzymes from a database for a given orphan reaction. We constructed the test set, named “Orphan-194”, comprising 194 orphan reactions. An “orphan reaction” refers to a reaction for which no catalyzing enzyme was recorded in databases before 2018 but was later annotated with enzymes after 2023. We provide more details on the construction of Orphan-194 in the Methods section.

EnzymeCAGE was benchmarked against several leading methods, including the similarity-based tool Selenzyme [40] and two deep learning-based methods: ESP [24], CLIPZyme [34]. For candidate enzyme selection, we computed the similarity between the target orphan reaction and all reactions in the training set, selecting the top 10 most similar reactions. The enzymes associated with these reactions, all discovered before 2018, were then used as candidates for the target orphan reaction when performing the retrieval task. Experimental results demonstrated that EnzymeCAGE can accurately retrieve enzymes for orphan reactions (Fig. 3). EnzymeCAGE showed a stronger ability to distinguish positive from negative enzyme-reaction pairs compared to baseline models, achieving higher binary classification AUC (Fig. 3b) and improved EF scores (Fig. 3c). To further analyze retrieval capability, we examined test set reactions with decreasing similarity thresholds to the training set (from 0.9 to 0.6). EnzymeCAGE consistently outperformed baselines across thresholds, achieving better Top-10 DCG results (Fig. 3d) and Top-k SR (Fig. 3e). Additionally, two visual examples are provided (Fig. 3f) to illustrate EnzymeCAGE’s effectiveness in retrieving compatible enzymes given reactions absent from the training set. These examples simulate the use case of applying EnzymeCAGE to orphan reactions.

**Fig. 3.**
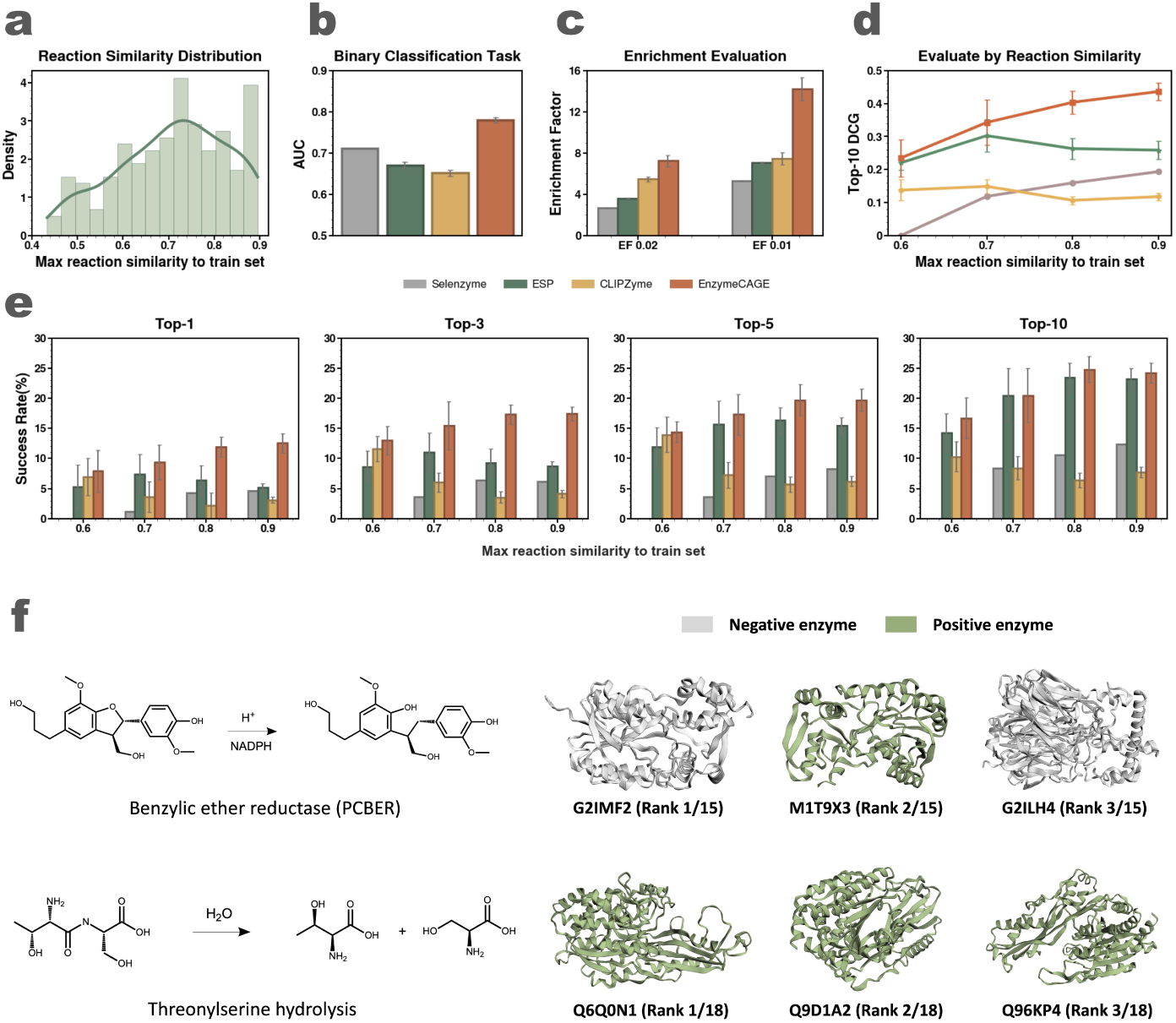
**EnzymeCAGE’s performance on test set Orphan-194**. **(a)** Data distribution of reactions within the Orphan-194 test set. **(b)** Prediction performance on the binary classification task. **(c)** Enrichment evaluation at ratios of 0.01 and 0.02. **(d)** Top-10 DCG evaluation across varying thresholds of maximum reaction similarity to the training set. **(e)**Top-k SR evaluation across different reaction similarity thresholds. **(f)**Examples of high-ranking predictions. Each row displays a chemical reaction on the left, with the top three ranked enzymes predicted by EnzymeCAGE on the right—positive enzymes highlighted in green and negative ones in yellow. In the first example, EnzymeCAGE identified benzylic ether reductase among 15 recalled enzymes, successfully ranking the positive enzyme in the 2nd position. In the second example, a threonylserine hydrolysis reaction, EnzymeCAGE retrieved 18 enzymes, ranking all positive enzymes within the top three positions.

### 2.4 Domain-specific Fine-tuning and EnzymeCAGE Performance on External Test Sets

Enzyme families in nature exhibit diverse functional and structural characteristics, with some families, such as Cytochromes P450, drawing considerable attention due to their important roles in biochemical processes and relevance to biotechnology and medicine. To enhance the predictive accuracy for these enzyme families, EnzymeCAGE incorporates domain-specific finetuning, allowing the model to adapt to the nuanced functional requirements within specific families. This approach enables targeted function prediction and improves performance in scenarios requiring domain knowledge.

To evaluate the effectiveness of domain-specific fine-tuning, we constructed external test sets for three prominent enzyme families, including Cytochromes P450 [54], terpene synthases [55], and phosphatases [56]. Fine-tuning starts with building a comprehensive training set tailored to the target enzyme family. This involves identifying all enzymes and their corresponding reactions within the family from the training data and generating all possible enzyme-reaction pairs. Post fine-tuning, EnzymeCAGE demonstrated marked improvements in performance (Fig. 4). For Cytochromes P450 and terpene synthases, fine-tuning notably increased the Top-k SR and reduced result variance, indicating enhanced model stability (Fig. 4a, 4c, 4e). In the case of phosphatases, where the initial performance of EnzymeCAGE fell short of baseline methods, fine-tuning brought the model’s performance to levels comparable to or surpassing baseline results. For each enzyme family, we selected two representative cases with favorable prediction outcomes (Fig. 4b, 4d, 4f). These evaluations on external test sets highlight that domain-specific fine-tuning empowers EnzymeCAGE to better capture family-specific characteristics, ultimately leading to more accurate, consistent, and reliable function predictions.

**Fig. 4.**
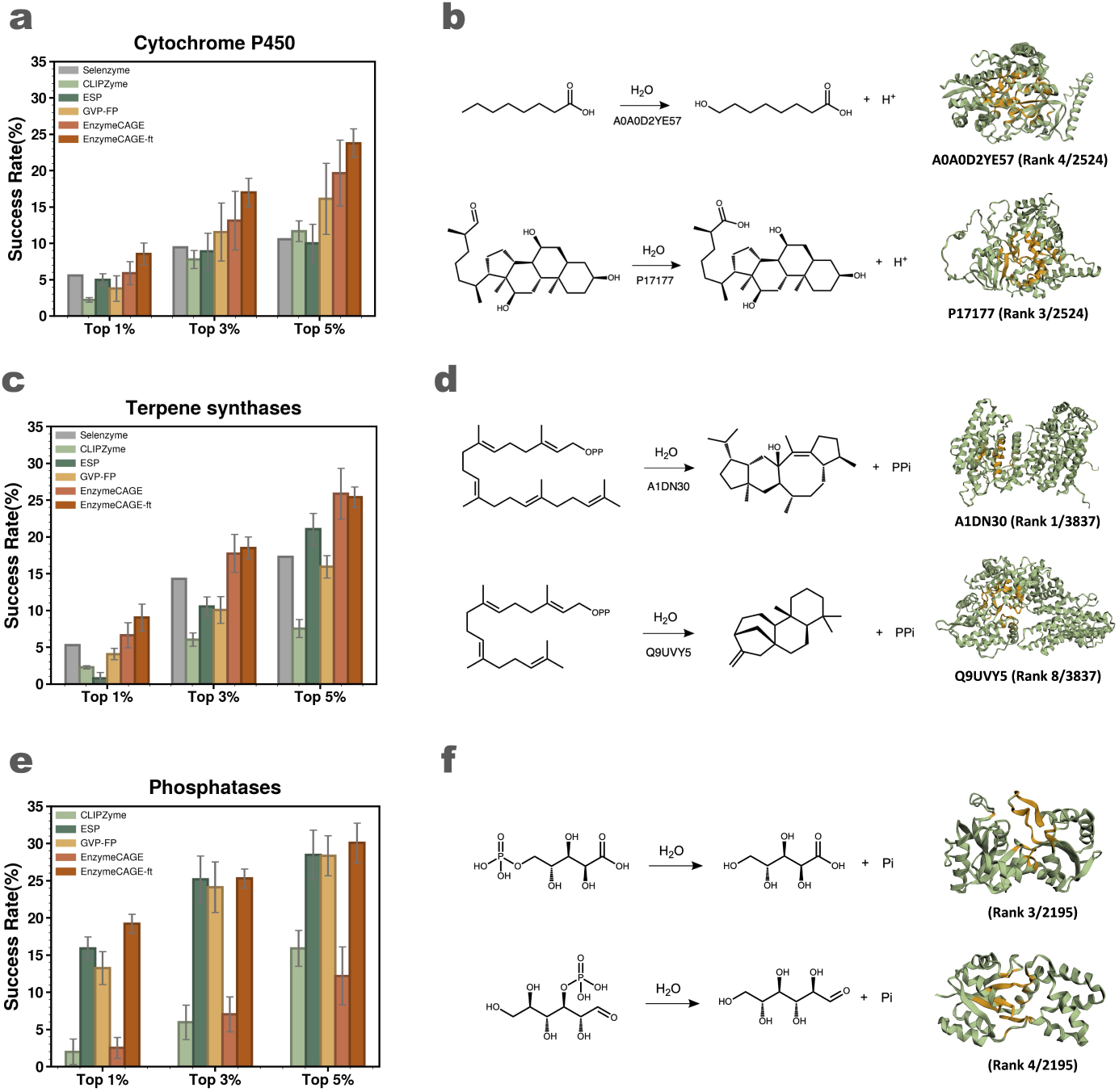
**EnzymeCAGE’s performance on external test sets**. **(a, c, e)** Top k% Success Rate (SR) evaluation on P450s, terpene synthases, and phosphatases datasets. “EnzymeCAGE-ft” indicates the model after domain-specific fine-tuning. **(b)**Examples of well-predicted P450-catalyzed reactions for two distinct scaffold substrates: In the first reaction, the hydrocarbon chain of a short-chain carboxylic acid undergoes terminal oxidation, with EnzymeCAGE ranking enzyme A0A0D2YE57 in 4th position out of 2,524 candidates, demonstrating its ability to identify active enzymes within the top 0.2%. In the second example, a terminal aldehyde group on a steroid scaffold is oxidized to a carboxyl group, with enzyme P17177 ranked 3rd, highlighting the model’s precision in pinpointing high-ranking enzymes. **(d)** Examples of well-predicted terpene synthase-catalyzed reactions for substrates GFPP and GGPP: in the first case, GFPP serves as the substrate, yielding a tetracyclic terpene compound with enzyme A1DN30 ranked first by EnzymeCAGE; in the second case, GGPP is the substrate, producing a tricyclic terpene product with enzyme Q9UVY5 placed within the top ten. **(f)** Two well-predicted examples of phosphatase-catalyzed hydrolysis reactions involving phosphate compounds.

### 2.5 EnzymeCAGE effectively retrieves suitable enzymes for biosynthetic pathways

One of the key strengths of EnzymeCAGE is its ability to facilitate the discovery and reconstruction of important molecular biosynthesis through enzyme retrieval. Glutarate, an important metabolic intermediate, has applications spanning food, pharmaceuticals, chemicals, agriculture, environmental remediation, and other domains. We selected a glutarate biosynthesis pathway [57] for a case study evaluating EnzymeCAGE’s enzyme retrieval performance.

This pathway consists of six reactions, starting from *α*-ketoglutarate, a key intermediate of the TCA cycle (Fig. 5a). To simulate real-world applicability, we removed all pathway-related reactions from the training set and re-trained EnzymeCAGE on the de-duplicated dataset. For each pathway reaction, we compared EnzymeCAGE’s enzyme retrieval performance with that of Selenzyme. Specifically, for each reaction, we selected similar reactions from the database with a similarity score above 0.6, then collected the corresponding enzymes for these reactions as candidates. EnzymeCAGE and Selenzyme were then used to score and rank these enzyme candidates.

**Fig. 5.**
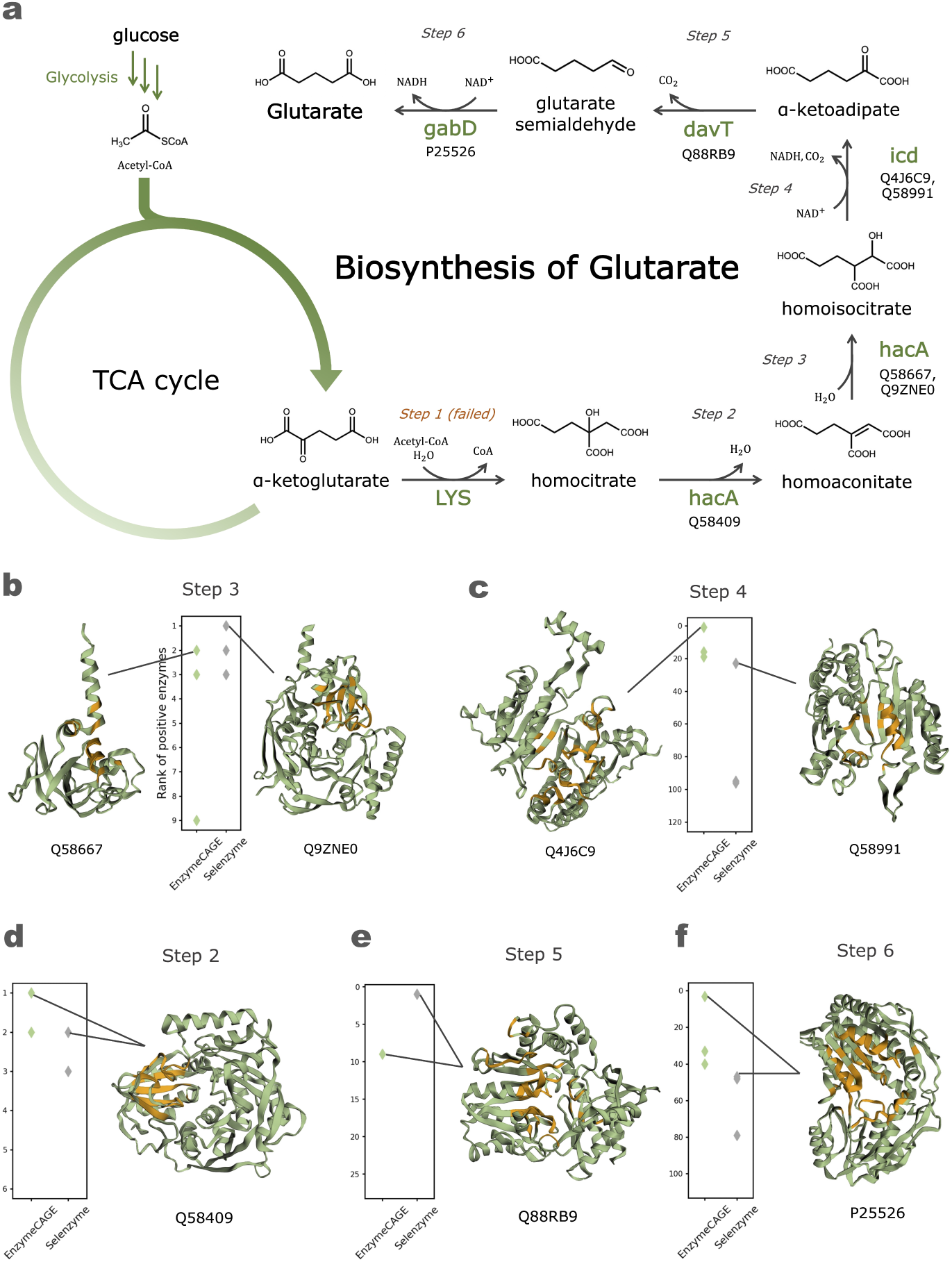
**Enzyme retrieval by EnzymeCAGE in the glutarate biosynthesis pathway**. **(a)** Diagram of the synthetic glutarate biosynthesis pathway, comprising six reaction steps. Green text beneath each reaction arrow represents the gene encoding the enzyme responsible for catalyzing the respective reaction, with the UniProts of successfully retrieved enzymes listed below each gene. **(b, c, d, e, f)** Enzyme retrieval results for each reaction step, illustrated with specific details for Steps 2 and 3. For Step 2, retrieval results are displayed in Fig. 5d, showing a scatter plot of the ranking positions of positive enzymes among the candidate pool. Here, EnzymeCAGE and Selenzyme identified 6 candidates, including 2 positive enzymes. The top-ranking positive enzyme for both methods is Q58409. For Step 3, retrieval results appear in Fig. 5b, where 9 candidate enzymes, including 3 positives, were evaluated. EnzymeCAGE’s highest-ranking positive enzyme is Q58667 (ranked 2nd), whereas Selenzyme ranked Q9ZNE0 as the top positive enzyme (ranked 1st).

In this evaluation, EnzymeCAGE ranked positive enzymes within the top 10 for all pathway reactions except Step 1, where no known positive enzyme was found. Notably, for Steps 2 (Fig. 5d), 3 (Fig. 5b), 4 (Fig. 5c), and 6 (Fig. 5f), EnzymeCAGE ranked positive enzymes within the top 3. In contrast, Selenzyme ranked positive enzymes well only for Steps 2, 3, and 5, each having fewer than 30 candidates. For Steps 4 and 6, where over 100 candidate enzyme sequences were present, EnzymeCAGE’s best-ranked positive enzymes were within the top 3, while Selenzyme’s were ranked beyond 20. This case study exemplifies EnzymeCAGE’s strong enzyme retrieval capabilities in complex biosynthetic pathways, highlighting its advantage over the de facto standard enzyme function annotation tool in pathway-specific applications [58, 59].

## 3 Discussion

Here we present a catalytic-specific structure-informed deep learning model for enzyme function prediction. EnzymeCAGE achieves state-of-the-art performance across two internal test sets (unseen enzymes and orphan reactions) and three external test sets (Cytochromes P450, terpene synthases and phosphatases). EnzymeCAGE not only successfully predicts which unseen enzymes correspond to known reactions but also accurately retrieves compatible enzymes for orphan (unseen) reactions, demonstrating high generalization capability in two directions. In the model design, EnzymeCAGE benefits from geometric guidance, embedding key information from both the enzyme and reaction into the model, which allows for better learning of the spatial structures and interactions between enzymes and reactions.

In traditional enzyme function prediction, methods typically encode the entire enzyme, including regions that may be functionally less relevant. This redundant information can make it challenging for models to learn enzyme functions accurately. A key advantage of our approach lies in the focused extraction of the enzyme pocket, focusing on relevant structural information and hypothetical pocket-specific enzyme-reaction interactions. Furthermore, the AlphaFill-predicted pockets do not rely on ground truth data for catalytic active sites or strictly defined pocket boundaries. As a result, the model’s predictions will exhibit a degree of robustness to minor errors or variations in pocket definitions.

For reaction encoding, we calculate a reacting area weight matrix as a form of geometric guidance to help the model more effectively capture and learn the reaction center. This involves determining atom-atom mappings to identify reactive sites. Accurately mapping reaction centers remains a significant challenge, particularly for highly complex biological reactions, due to the lack of universally reliable ”ground truth” atom mapping information. To address this limitation, the reacting area weight matrix is designed to be flexibly applied as optional geometric guidance, serving as attention weight adjustments. This allows the model to leverage this additional information when available, without making it a strict requirement for all data points. To further enhance model performance, future work could focus on developing more robust atom-atom mapping tools for large, complex reactions. This advancement would improve reaction center extraction, leading to better data quality and, ultimately, more effective enzyme function modeling.

In our ablation experiments, we systematically assessed the contribution of each module in EnzymeCAGE (Fig. 2g). The results clearly demonstrate that each component is essential for enhancing the model’s performance. Specifically, the inclusion of DRFP for reaction representation is critical to the model’s success. Removing DRFP led to a significant drop in performance, underscoring its importance in effectively capturing the reaction space. Furthermore, incorporating both the catalytic pocket and the molecular 3D structure from the reaction proved to be crucial. When these structural elements were excluded, the model relied solely on the ESM embeddings and DRFP, which resulted in the poorest performance across all ablation conditions. These findings reinforce the overall design of EnzymeCAGE, highlighting that each module plays a vital role in improving its predictive capability.

In the data construction process, a major challenge is the lack of negative samples of enzyme-reaction pairs. To solve this problem, we constructed negative samples by considering both negative reactions and negative enzymes, balancing the difficulty of distinguishing between positive and negative samples with the rationality of negative data. However, the presence of false-negative data samples cannot be completely avoided. A potential solution to this issue is better collection and labeling of negative samples, especially for data that diverge from “real-world” expectations. These data are important for reaction modeling and function prediction, enabling the model to better capture the effects of subtle structural differences on catalytic activity.

Although EnzymeCAGE primarily focuses on enzyme retrieval and function prediction by predicting enzyme-reaction interactions, the model also provides interpretability in its predictions. When using the geometric crossattention model to learn the interaction between enzyme and reaction, we extract attention weights and use them to identify active sites. In many enzymes, this method accurately predicts the location of active sites. However, a current limitation is that we can only identify active sites located within the enzyme pocket as estimated by AlphaFill, while in reality, some active sites may lie outside the pocket but are involved in catalysis.

## 4 Methods

### 4.1 Data Construction

To build a robust dataset for enzyme-reaction prediction, we collected curated and validated enzyme-reaction pairs from RHEA [14], MetaCyc [15], and BRENDA [47]. We optimized the dataset by removing invalid entries and standardizing formats. Specifically, enzyme entries without UniProt ID [13] or sequence data were excluded. For reactions, we applied the following criteria: (1) remove cofactors, ions, and small molecules present on both substrate and product sides, (2) exclude reactions involving more than five substrates or products, and (3) standardize molecular SMILES using OpenBabel [60]. This refinement yielded 328,192 enzyme-reaction pairs, comprising 145,782 unique enzymes and 17,868 distinct reactions.

To enhance model training, we generated negative samples. We created two types of negatives: (1) Negative Enzymes for Reactions – Given a reaction, we identified catalyzing enzymes and retrieved homologous proteins and dissimilar enzymes (in a 2:1 ratio) as negative examples(Supplementary Fig. S3). Homologous proteins with greater than 40% sequence similarity were identified using MMseqs2 [61]. Dissimilar enzymes were selected by choosing enzymes with different top-three-level EC classifications. (2) Negative Reactions for Enzymes – We used RXNMapper and RDChiral [62] to generate reaction templates with atom-atom mappings and EC annotations. For each positive enzyme-reaction pair, we applied a non-corresponding reaction template to the enzyme’s substrate to generate a new product and negative sample. This augmentation process expanded the training set to 914,077 enzyme-reaction pairs.

We constructed two internal test sets, Orphan-194 and Loyal-1968, for benchmarking on orphan reactions and unseen enzymes. For the test set Orphan-194, we first filtered the full dataset to retain only reactions with EC numbers found in the training set and identified 690 reactions without EC numbers as candidates for the test set. These reactions have a similarity score below 0.9 compared to any reaction in the training set and are catalyzed by at least five enzymes. We then applied a temporal filter, selecting only reactions that had no recorded catalyzing enzyme in the RHEA database before 2018 but were later annotated with enzymes after 2023. This refinement yielded a final test set of 194 reactions, designated as Orphan-194.

For the Loyal-1968 test set, we identified non-promiscuous enzymes that have been reported to catalyze only a single reaction. Using MMseqs2 with a cutoff of 0.4, we retrieved homologous proteins as negative samples. This set was curated to include reactions with: (1) at least one non-promiscuous enzyme as a positive example, (2) each non-promiscuous enzyme having over 10 homologous proteins, and (3) a ratio of homologous to non-promiscuous enzymes of at least 3:1. These criteria ensure functional diversity and exclude positive enzymes present in the training data. The final Loyal-1968 test set consists of 455 reactions with a total of 71,853 enzyme-reaction pairs, covering 1,968 enzymes across 77 unique top-three-level EC numbers.

### 4.2 Catalytic Pocket and Reaction Center Extraction

To generate catalytic pocket data, we downloaded AlphaFold structures for all enzymes and applied AlphaFill [44] to extract the enzyme pockets. For identifying reaction center, we used atom-atom mapping of the reactions. In the pocket extraction process, AlphaFill first identified homologous proteins of the target enzyme in the PDB-REDO database [63] and located their protein-ligand complexes. Ligands from these homologous complexes were then transplanted onto the target enzyme structure through structural alignment (Fig. 1a). Following ligand transplantation, we selected ligands based on their atomic count and occurrence frequency and defined the catalytic pocket using an 8 Å radius around each ligand. A clustering analysis on the extracted pockets using Foldseek [64] showed that catalytic pockets have higher functional relevance than entire enzyme structures, supporting the focus on the pocket-level analysis and modeling (Supplementary Fig. S1). To identify reaction center, we employed RXNMapper to generate atom-atom mappings between substrates and products. Using this mapping, we identified atoms involved in bond changes, charge shifts, and chirality alterations, marking these as the reaction center for the catalyzed transformation.

### 4.3 EnzymeCAGE Model and Architecture

EnzymeCAGE takes both the enzyme structure and the target reaction as inputs, starting by identifying the catalytic pocket and reaction center. The catalytic pocket is represented as a graph, where each node represents a residue enriched with features such as residue type, dihedral angles, orientations, and side-chain information. Each edge encodes spatial connections between residues, incorporating relative positional vectors and distance encodings (with distance encoded via radial basis functions). EnzymeCAGE employs a graph neural network, specifically GVP [65], to encode these pocket graphs due to its computational efficiency. Beyond encoding catalytic pocket features locally, EnzymeCAGE uses ESM2 to capture global features at the full-enzyme scale, thereby integrating structural and sequential information at both local and global levels. To process reactions, EnzymeCAGE begins by computing substrate and product conformations. It identifies the reaction center by pinpointing atoms that undergo changes in bonding, charge, or chirality during catalysis. Different weights are then assigned to atoms within the reacting area, with higher weights for atoms located in the reaction center. SchNet [66] is used to separately encode substrate and product conformations, providing an effective representation of reaction dynamics.

#### 4.3.1 Geometry-enhanced Pocket Attention Module

This module facilitates interaction between residues within the catalytic pocket by representing the pocket as a graph, *G_e_* = (*V_e_, E_e_*). Here, *V_e_* denotes node features such as residue types, residue positions, dihedral angles, orientations, and side-chain information, while *E_e_* represents edge features, including relative positional vectors and RBF-encoded distances. To further refine residue relationships, a residue-level distance map *D_m_* = {*d_ij_* | *i, j* ∈ *V* } is constructed, where *d_ij_* represents the distance between residues *i* and *j*.

Using this distance map, a geometry-enhanced attention mechanism captures local interactions within the pocket:

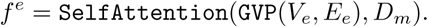

Here, a multi-head self-attention model [67] is applied, with the distance map *D_m_* serving as an attention bias to enhance the model’s understanding of relative positional relationships among residues. The resulting output, *f^e^*, encodes the local geometric and biochemical information of the catalytic pocket.

#### 4.3.2 Center-aware Reaction Interaction Module

This module captures the dynamic transformations between substrate and product molecules in a reaction, generating a comprehensive reaction representation. First, for each molecule (either substrate or product), we define a set *C* of indices corresponding to atoms at the reaction center. We then create a vector *R* to indicate these centers and calculate a reacting area weight matrix as follows:

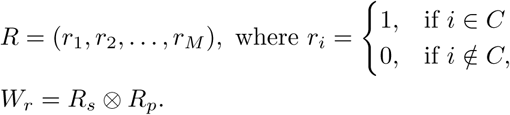

Here, *r_i_* represents an element of the vector *R*, and *M* is the total number of atoms in the molecule. *R_s_* and *R_p_* denote the reaction center of the substrate and product, respectively, and *W_r_* is the reacting area weight matrix for the reaction. This matrix will guide the reaction interaction model by emphasizing reaction center.

The substrate and product conformations are represented as graphs, *G_s_* = (*V_s_, E_s_*) and *G_p_* = (*V_p_, E_p_*), respectively, where each node represents an atom with features like atom type and position, and each edge represents a bond between atoms. A 3D molecular GNN, specifically SchNet, encodes these substrate and product graphs:

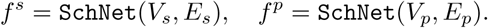

Using the molecular embeddings *f^s^* and *f^p^*, we build an interaction module based on cross-attention, where the reacting area weight matrix *W_r_* and molecular representations serve as inputs:

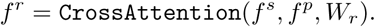

In this module, the reacting are weight *W_r_* acts as an attention bias, guiding the model to focus effectively on the reaction center. The resulting output, *f_r_*, represents the complete reaction.

#### 4.3.3 Geometry-enhanced Interaction Module

This module models the interactions between the catalytic pocket and the chemical reaction, employing geometric guidance to determine enzyme-reaction interaction weights. Given that the active site of an enzyme resides within the catalytic pocket (where the active site is a subset of catalytic pocket residues), we start by calculating a conservation score for each residue based on its proximity to the pocket center:

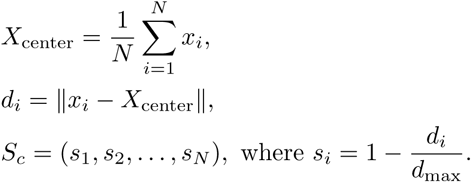

Here, *X*_center_ represents the geometric center of the pocket, *N* denotes the number of residues, and *S_c_* is the conservation score vector, where residues closer to the pocket center receive higher scores.

Next, we compute the enzyme-reaction interaction weight matrix by combining the pocket conservation scores with the reaction center weights of the substrate:

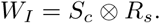

In this expression, *W_I_*denotes the enzyme-reaction interaction weight matrix, which is used as a geometric guidance in the cross-attention model to enhance the enzyme-reaction interaction:

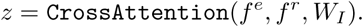

In this setup, *f^e^* and *f^r^* represent the feature embeddings of the catalytic pocket and the reaction, respectively, while *W_I_* provides the interaction weights based on geometric guidance. The output *z* is the final embedding produced by the Geometry-enhanced Interaction Module, capturing the interaction between the catalytic pocket and the chemical reaction(Supplementary Fig. S2).

#### 4.3.4 Global Feature Integration

To generate the final prediction, EnzymeCAGE combines local pocket-reaction embeddings, global enzyme embeddings, and reaction embeddings. EnzymeCAGE encodes the full enzyme using ESM2, producing a global enzyme representation denoted as *f*_esm_. For the reaction, EnzymeCAGE uses the reaction fingerprint DRFP, producing the reaction embedding *f*_DRFP_.

The global enzyme representation *f*_esm_ and the reaction fingerprint *f*_DRFP_ are concatenated with the local catalytic pocket embeddings *z*, and a multilayer perceptron (MLP) is applied to predict catalytic specificity:

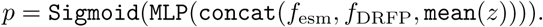

The final prediction *p* represents the probability that the input enzyme can catalyze its associated given reaction, offering a quantitative measure of catalytic specificity.

### 4.4 External Test Sets Construction

To evaluate the performance on external test sets, we selected three prominent enzyme families: Cytochromes P450, terpene synthases, and phosphatases. For each family, we curated and processed an open-source dataset to ensure highquality evaluation. For Cytochromes P450, we utilized P450Rdb, a manually curated dataset containing over 1,600 reactions involving approximately 600 P450 enzymes from more than 200 species. For terpene synthases, the dataset comprised over 400 relevant reactions and more than 1,100 terpene synthases. Lastly, for phosphatases, we employed a high-throughput screening dataset consisting of 165 reactions, 218 phosphatases, and 35,970 enzyme-reaction pairs. Each dataset was subjected to the same preprocessing pipeline as the internal test sets, including data cleaning and the exclusion of reactions with a similarity score exceeding 0.9 to those in the training set. This preprocessing step yielded three well-defined external test sets for evaluation.

### 4.5 Domain-specific Fine-tuning Details

In the domain-specific fine-tuning process, we began by constructing a comprehensive fine-tuning dataset for the target enzyme family. This involved identifying all enzymes and their associated reactions within the family from the training data and generating all possible enzyme-reaction pairs. Pairs already present in the original training data retained their original labels, while newly generated pairs were assigned as negative samples. This approach yielded training sets containing between 160,000 and 280,000 enzyme-reaction pairs for fine-tuning. Subsequently, we utilized the pre-trained model from the evaluation scenario associated with the Loyal-1968 test set as the base model and performed fine-tuning for a fixed number of five epochs for each enzyme family scenario.

### 4.6 Reaction Similarity Measure

Our approach for computing reaction similarity primarily referred to the method used in Selenzyme. The similarity between two reactions is computed by comparing their reactants and products using molecular fingerprints and the Tanimoto coefficient. Each molecule in a reaction is represented by a Morgan fingerprint with a radius of 8. The similarity between two molecular fingerprints *A* and *B* is quantified using the Tanimoto coefficient. For two reactions *R*_1_ and *R*_2_, we calculate two similarity scores: direct matching (*S*_1_) and cross matching (*S*_2_). *S*_1_ compares reactants with reactants and products with products:

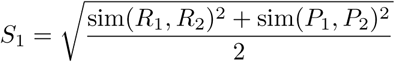

Where R represents the reactants and P represents the products. *S*_2_ compares reactants with products and vice versa:

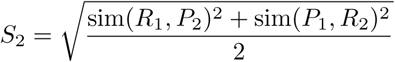

The final similarity score for the reaction pair is the maximum of *S*_1_ and *S*_2_. This approach ensures comprehensive comparisons between reactions, accounting for both direct and cross-reactivity relationships.

## Data availability

We used freely available data as described in Methods. Databases including RHEA (https://www.rhea-db.org), BRENDA (https://www.brenda-enzymes. org) and MetaCyc (https://metacyc.org) were used in training dataset construction. Databases including P450Rdb (https://www.cellknowledge. com.cn/p450rdb), Terpene Synthase data (https://zenodo.org/records/ 10567437) and Phosphatases data (https://github.com/samgoldman97/ enzyme-datasets) were used in external test sets construction. AlphaFold protein structure database (https://alphafold.ebi.ac.uk/) was used to obtain the 3D structure of the enzymes. PDB-REDO (https://pdb-redo.eu/) was used in the catalytic pocket extraction by AlphaFill.

## Code availability

The source code of EnzymeCAGE is available at https://github.com/ GENTEL-lab/EnzymeCAGE.git.

## Acknowledgements

We thank Wei Lu and Jixian Zhang for valuable discussions and feedback. We are grateful to Meihui Song for their visualization and inspiration. This study has been supported by the National Natural Science Foundation of China [62041209] and the Key project at CG level: The ability establishment of sustainable use for valuable Chinese medicine resources [2060302].

## Authors’ contributions

S.Z. conceived and supervised the project. Y.L. and J.Z. contributed to the algorithm implementation. Y.L., T. Z and R. W contributed to the data collection. Y.L.,C.H., J. R., C.C and S.Z. contributed to the visualization and server implementation. Y.L.,C.H. and S.Z. wrote the manuscript. All authors were involved in the discussion and proofread.

